# Neurons in the postrhinal cortex encode nonspatial context during visual biconditional discrimination

**DOI:** 10.1101/2022.06.23.497405

**Authors:** Victoria R. Heimer-McGinn, Sean G. Trettel, Brendon Kent, Maya A. Singh, Rebecca D. Burwell

## Abstract

Spatial context, or the physical surroundings that form the background of an experience, is an essential component of episodic memory. The rodent postrhinal cortex and its primate homolog, the parahippocampal cortex, are thought to preferentially process visuospatial information to represent the spatial features of contexts and scenes. In this study, we addressed open questions about postrhinal function and about how context modulates behavior and cognition. The first question was whether the postrhinal cortex also represents nonspatial contexts. The second question was how representations of context might interact with other cues in the environment. We recorded postrhinal neurons as rats performed a visual nonspatial biconditional discrimination task in which the pattern on the floor determined which object in a pair was correct. Critically, this task design allowed dissociation of location from non-spatial context. We found that postrhinal ensembles and neurons signaled changes in non-spatial context and coded for conjunctions of non-spatial context and objects. Importantly, postrhinal neurons coded for conjunctions of context and objects more often than they coded for conjunctions of location and object. The pattern of findings suggests that postrhinal representations of context may behave like occasion setters by modulating the meaning of other cues in the environment.

## Introduction

The rodent postrhinal cortex (POR) and its primate homolog, the parahippocampal cortex (PHC), preferentially process visuospatial information and provide such information to the hippocampus, whereas the perirhinal cortex (PER) provides nonspatial object or pattern information to the hippocampus. One view of hippocampal information processing is that spatial and nonspatial information arrive to the hippocampus by separate processing streams such that the spatial processing stream includes the postrhinal and medial entorhinal cortices and the nonspatial processing stream includes the perirhinal and lateral entorhinal cortices (Eichenbaum et al., 2012). The hippocampus is thought to bind spatial and nonspatial information in the service of episodic memory and associative learning. However, it is increasingly clear that these pathways are not well segregated. Although the POR and the PHC do project strongly to the medial entorhinal cortex, they are also robustly interconnected with the PER and lateral entorhinal cortex, regions associated with the so called nonspatial pathway (Burwell and Amaral, 1998, Suzuki and Amaral, 1994, Doan et al., 2019). Indeed, disconnection of the POR and the PER, which presumably leaves these processing streams intact, impairs recognition of objects in particular contexts (Heimer-McGinn et al., 2017). This suggests the POR has access to nonspatial object and item information arriving directly from the PER. One possibility is this robust combination of spatial and nonspatial information upstream of the hippocampus supports functions previously attributed to the hippocampus. Along these lines, we have suggested that the PER provides object and pattern information to the hippocampus for associative learning and episodic memory and to the POR for representing the local physical context (Estela-Pro and Burwell, 2022, Ho and Burwell, 2014, Peng and Burwell, 2021, Heimer-McGinn et al., 2017). POR representations of context would also be available to other brain regions for context-guided behavior and cognition, for example, the prefrontal cortex (Hwang et al., 2018, Kondo and Witter, 2014).

Evidence in humans, monkeys, and rodents shows that the rodent POR and its homolog, the primate PHC, play an important role in processing information related to spatial context including the geometry of spaces, landmark configurations, spatial layout of objects, and spatial object characteristics such as size and portability (Mullally and Maguire, 2011, Martin et al., 2013, LaChance et al., 2022, Bonner and Epstein, 2021, Bachevalier et al., 2015). Human fMRI studies show that the PHC is activated in response to scenes, landmarks, and objects in spatial context (Aminoff et al., 2013, Bar and Aminoff, 2003, Bonner and Epstein, 2021), and PHC damage impairs landmark identification, navigation, and spatial memory (Aguirre and D’Esposito, 1999, Ploner et al., 2000). Similarly, PHC lesions in monkeys impair both object-location and object-in-context recognition (Malkova and Mishkin, 2003, Bachevalier et al., 2015). In rodents, POR lesions impair context discrimination (Bucci et al., 2002), scene discrimination (Gaffan et al., 2004), object-in-context recognition (Norman and Eacott, 2005), and contextual fear conditioning (Bucci et al., 2000, Burwell et al., 2004a). POR lesions also impair contextually driven spatial choices (Park et al., 2017). Moreover, disconnection of the POR and PER results in object-in-context impairment, suggesting that POR relies on object and pattern information transmitted directly from PER to encode representations of context (Heimer-McGinn et al., 2017). Interestingly, the POR is also responsive to objects in motion (Nishio et al., 2018) and to changes in the local environment (Burwell and Hafeman, 2003). Finally, POR responds to both objects and locations and encodes object-location conjunctions (Furtak et al., 2012).

The present study was designed to address two open questions about the function of the POR. The first question is whether the POR has a role in representing non-spatial contexts. There are some hints of this possibility in the literature. Diana (2017) reported that the PHC is activated by context retrieval in a source memory paradigm that minimized spatial processing by crafting encoding questions that drew upon lexical characteristics of the words to be remembered. Additionally, fMRI studies in humans show that the PHC responds to objects with strong contextual associations, for example, the target word “microscope” is strongly associated with laboratories (Bar and Aminoff, 2003).

The second open question is how POR representations of context might modulate behavior. One theory is that context operates as an occasion setter (Fraser and Holland, 2019). Occasion setting refers to the capability of a stimulus to modulate the meaning of other cues, for example, the association of another stimulus with a reward. Thus, stimuli can be directly associated with one another as in stimulus-stimulus associations, or a stimulus can modulate the meaning of another stimulus by acting as an occasion setter. Stimuli tend to operate as occasion setters when separated from other stimuli by longer intervals and when the occasion setter differs from other stimuli in character or modality. This perspective is particularly useful in understanding how nonspatial contexts might guide behavior.

To address our two open questions, we developed a nonspatial contextual biconditional discrimination (nscBCD) task in which a non-spatial context determined which of two objects was correct. The task was designed to favor occasion setting in two ways. The non-spatial context differed from the objects in character, and the presentation of the non-spatial context was separated in time from the presentation of the objects. In the nscBCD task, rats learned to discriminate which of two 2-dimensional (2D) objects presented on the floor would be rewarded based on a nonspatial context operationalized as the pattern on the floor. The task was conducted in a bow-tie shaped maze, such that presentation of pairs of objects alternated from one side of the maze to the other. One of two floor patterns was randomly selected at the beginning of each trial. Because objects were presented in two locations, we were able to dissociate neural correlates of non-spatial context from neural correlates of location. Neural recordings collected during task performance were analyzed to show neural correlates of context onset as well as object stimulus onset. We also identified selectivity for conjunctions of objects, locations and contexts during task relevant epochs. To determine how objects, locations and contexts were represented by the ensemble of recorded single units, we used population similarity analyses. Our results provide evidence for the hypothesis that the POR represents non-spatial contexts and that those representations are used to guide behavior in a manner consistent with occasion setting.

## Results

Subjects were shaped and trained on the novel nscBCD task in which two 2D object stimuli were back-projected onto the floor of a bowtie-shaped maze in an automated manner (Figures 1A, 1B). The pattern on the floor determined which of a pair of objects was correct. There were two pairs of objects and two different floor patterns (Figure 1C). For each pair of objects, one was correct on the solid floor and the other was correct on the striped floor regardless of allocentric location (east or west) or egocentric location (left or right). One of two background patterns was randomly chosen for the next trial. Thus, sometimes the floor was the same as the prior trial, and sometimes the floor was changed. At the beginning of each trial, the onset of white noise signaled the rat to enter the ‘ready position’ in the center of the maze (Figure 1D). Trials alternated between east and west. Correct choices were counterbalanced for egocentric location (left/right) such that the correct choice could be presented in one of four possible locations (east left, east right, west left, and west right). Stimuli were presented when rats remained in the ready position for a variable fixation period (0.8 - 1.8 sec). A choice was made when the rat approached one of the two objects. A correct choice was rewarded, whereas an incorrect choice terminated the trial and no reward was delivered. Overall, there were 32 trial types (4 locations, 4 objects, and 2 floor patterns). Rewards were given in the form of medial forebrain intracranial stimulation (ICS) consisting of bipolar electrical pulses delivered to the medial forebrain bundle. The 100 Hz ICS pulses were 500 μsec in length, separated by 500 μsec, and sustained for 500ms. Amplitude varied depending on the sensitivity of the subject.

**Figure 1.**
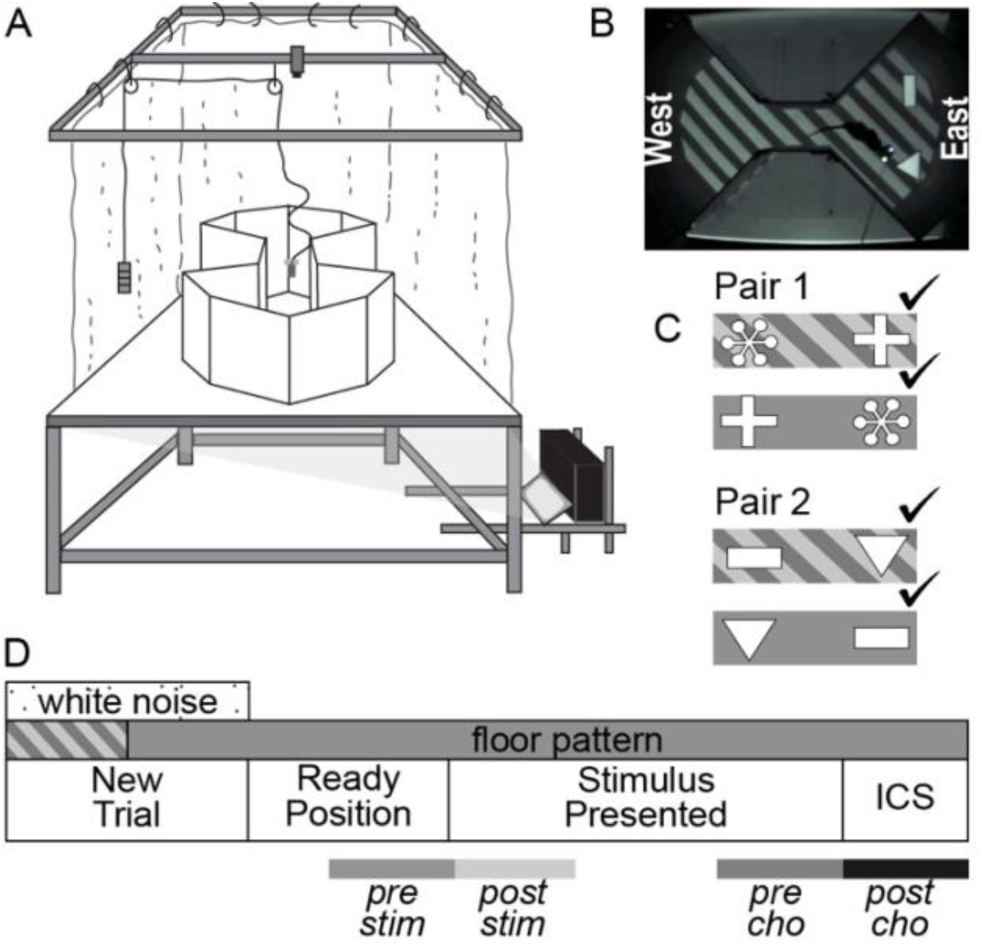
Behavioral Task. Overview of the behavioral task used in this study. (A) Schematic of the Floor Projection Maze. (B) Overhead view of a rat in the maze. (C) For each of the four possible discrimination combinations, the correct object-floor pairing is indicated by a checkmark. Each combination could appear on either the west or east side of the maze, the correct object could appear on the egocentric left or right, and the objects could be presented on either of two floor patterns. This yielded 32 trial types. (D) At the start of each trial, the context was randomly assigned. White noise signaled the rat to go to the ready position in the center of the maze and maintain the ready position for a variable time (800-1800 ms), after which the object pair was presented. The rat made a choice by approaching an object. The choice was recorded and both stimuli disappeared. Correct choices were rewarded with intracranial stimulation (ICS) to the medial forebrain bundle. Analysis epochs (500 ms) are represented by gray boxes at the bottom of the panel: pre-stim = *pre-stimulus* epoch, post-stim = *stimulus* epoch; pre cho = *pre-choice* epoch; post cho = *post-choice* epoch.

### Task Performance

Rats were first trained on a series of shaping steps (Supplemental Table S1). Upon reaching a criterion of 10 correct choices in 12 consecutive trials, rats progressed to the next shaping step. The mean number of sessions required for each stage was as follows: hand shaping, 5; auto shaping, 5; simple discrimination, 3.5; biconditional discrimination training with the first pair, 6.5; biconditional discrimination with the second pair, 6.5; and 1.5 days on the full nscBCD task with both pairs. Once trained, rats performed daily sessions of 150-200 trials, and sessions were analyzed when performance was above 55% accuracy.

### Histology and Unit Isolation

Stereotrodes were implanted in the POR of five animals. We recorded and isolated 120 cells from 21 stereotrodes across 54 sessions. Recordings spanned the POR along the rostro-caudal axis from −8.40 mm to –9.48 mm from Bregma (Figure 2). Stereotrodes were lowered ∼20 μm after each daily session, and thus new neurons were recorded in each session. Recorded units were sorted using a variety of manual and partially automated techniques for isolation based on waveform characteristics in Offline Sorter (Plexon, Inc. Dallas, TX). Only well-isolated units with a signal-to-noise ratio greater than 2:1, and spike separation of at least 1 ms were retained for further analysis. At the end of training, electrode location was marked by electrolytic lesion, rats were euthanized, and the brains were processed to identify the location of electrode tips.

**Figure 2.**
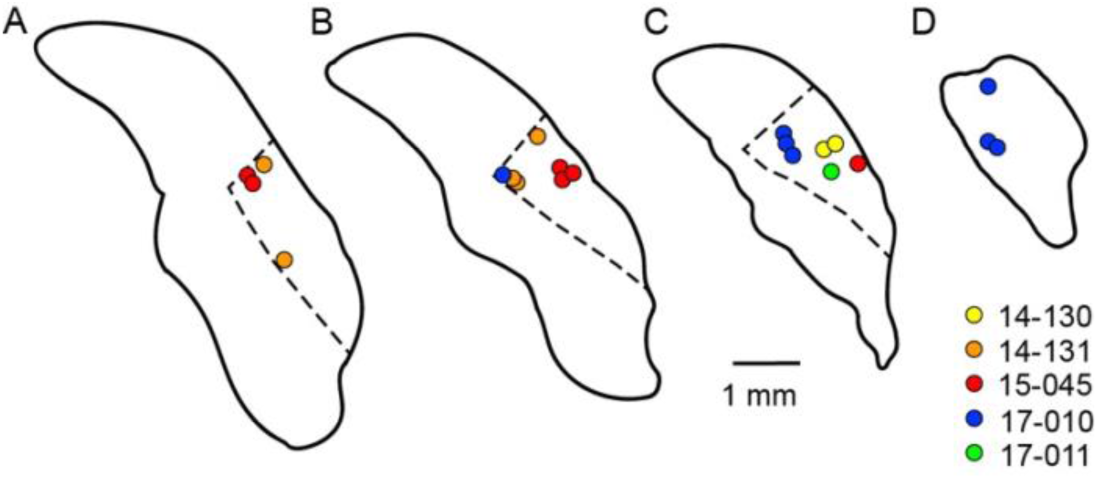
Histology. Schematic drawings show the location in the POR of lesions marking the tip of the electrode at the end of recording, color coded by animal. Contours are displayed in the coronal plane at −8.40 (A), −8.72 (B), −9.24 (C) and –9.48 (D) mm from Bregma. Dotted lines indicate POR boundaries.

### POR neurons signal changes in non-spatial context

To determine whether the POR signals a change in non-spatial context, we analyzed firing rate (FR) before and after context (floor pattern) changes. The goal of this analysis was to determine whether POR cells were selective for the onset of a new floor and/or the onset of a particular floor. For the 120 isolated cells, the mean baseline FR was 1.95 Hz and ranged from 0.2 to 12.94 Hz. A total of 104 cells met criterion for this analysis: at least 20 trials in which the floor changed from gray to stripe or vice versa, and at least 20 spikes in the 500 ms before (*prefloor)* and after (*postfloor)* floor onset. For each cell, we conducted a 2 X 2 repeated measures analysis of variance (rANOVA) with ‘Floor Change’ as the within-trial factor (*prefloor* vs *postfloor*), ‘Pattern’ as the between-trial factor (stripe-to-gray or gray-to-stripe), and FR as the dependent variable. We predicted that POR cells would change FR in response to a change in floor pattern (main effect of ‘Floor Change’) and that the magnitude and/or direction of that change may vary depending on the direction of the floor change, either stripe-to-gray or gray-to-stripe (‘Floor Change x Pattern’ interaction). When cells were selective for both the change of the floor and the change to a particular floor, we interpreted only the interaction.

Based on our hypothesis, we expected the POR to signal a change in context, i.e. the switch to a different floor pattern. We found that 50% of recorded POR cells signaled a change in floor pattern (Table 1). Twenty-six percent showed a main effect of Floor Change indicating that they signaled a change in the floor, irrespective of which floor appeared (Figure 3A). This finding is consistent with prior evidence that the POR monitors the current context for changes. Of particular interest, however, were the 24% of POR cells that signaled the onset of a particular floor pattern. For example, one cell increased its FR when the gray floor was switched to the striped floor (Figure 3B; left panel) but not on striped to gray changes (Figure 3B; right panel). These cells, which were identified by significant interactions of ‘Floor Change x Pattern’ (*p* < 0.01), provide evidence that the POR signals changes in non-spatial context. The remaining 50% of cells (52 out of 104) did not show selectivity for the change in floor pattern. Since floor pattern serves as a non-spatial context and guides behavior in this task, our results provide further evidence that POR neurons represent features of the environment that may serve as a context to guide behavior.

**Figure 3.**
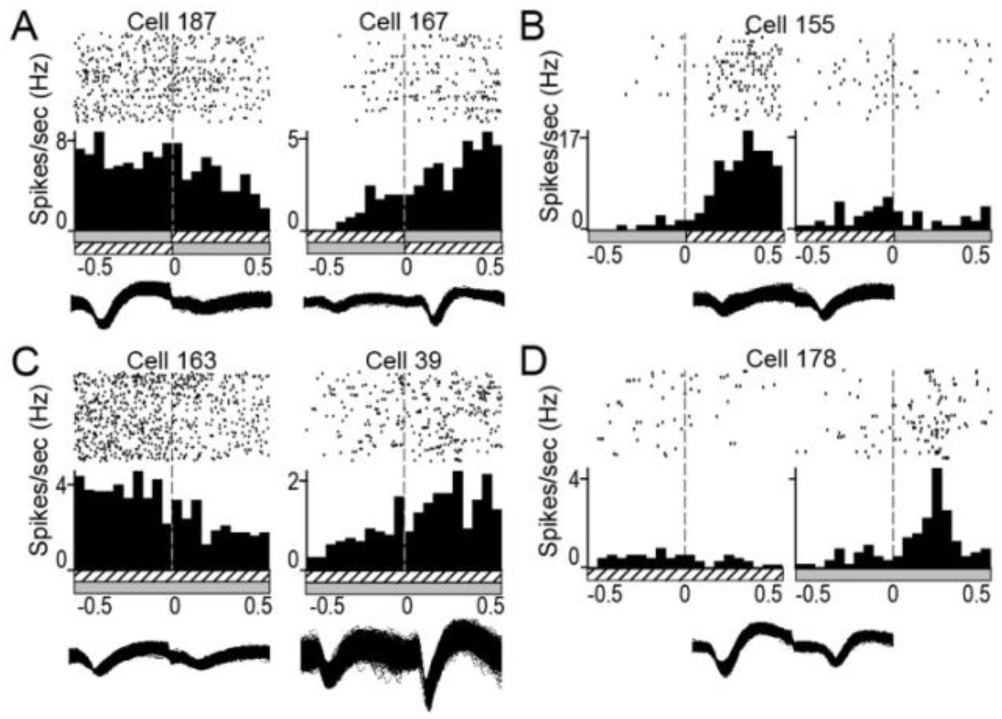
POR neurons signal changes in non-spatial context and the onset of 2D objects. Raster plots, perievent histograms, and wave forms are shown for representative cells displaying behavioral correlates. Postrhinal cells exhibit neuronal correlates during both floor change (A-B) and stimulus onset (C-D). (A) Examples of cells that increase or decrease firing rate (FR) in response to a change in floor pattern, irrespective of the identity of the floor. (B) An example cell that increases FR when the floor changes from gray to striped but not when it changes from striped to gray. (C) Examples of cells that increase or decrease FR when object stimuli appear, irrespective of object or floor identity. (D) An example of a cell that increases FR in response to object onset, but only when the context is a gray floor. Time bins = 25ms. Dashed lines represent floor or stimulus onset.

**Table 1.** POR neurons signal context change.

| Significant main effects and interactions | Number of cells | Percent |
| --- | --- | --- |
| Floor Change | 27 | 26 |
| Floor Change*Pattern | 25 | 24 |
| No response | 52 | 50 |
| **TOTAL** | **104** | **100** |

Of the 25 Floor Change\*Pattern cells, 11 also had a main effect of Floor Change.

### POR neurons signal the presentation of objects

We have proposed that the POR represents the spatial layout of features of the current context and monitors the current context for changes. Accordingly, we predicted that the POR would signal the presentation of objects on the floor and would represent the location of particular objects. Additionally, if the floor pattern was functioning as a context, the POR should signal the presentation of particular floor patterns and object pairs in the form of object-context conjunctive coding. *Importantly, in this analysis we used object pairs as opposed to individual objects because at this point in the task, a pair of objects had been presented, but an object had not yet been chosen*.

To determine the extent to which POR cells signaled the onset of object pairs, we analyzed the change in FR before and after the onset of 2D stimuli. Of the 120 cells we recorded and isolated, 114 met criteria for this analysis. The criteria were at least twenty trials for each of the context-object pairs (striped-pair1, striped-pair2, gray-pair1, and gray-pair2) and at least 20 spikes in the 500 ms before and after object stimulus onset. For each cell, a repeated measures ANOVA was carried out with ‘Stimulus Onset’ as the within-trial variable (*500 ms* before *vs 500 ms* after). Between-trial variables were ‘Context’ (gray or striped) and object ‘Pair’ (pair 1 or 2). We considered main effects to represent simple, non-conjunctive coding or selectivity and interactions to represent conjunctive coding or selectivity. We predicted that the FRs of POR cells would change in response to the onset of the object stimuli, and that this might, in some cases, depend on either the Context or the Pair.

We found that 44 of 114 cells (38.6%) signaled the onset of object stimuli either by a main effect of Stimulus Onset, an interaction with Stimulus Onset, or multiple main effects and/or interactions (Table 2). Some POR cells (25 of 114; 22%) responded to the object stimulus onset regardless of floor identity or object pair by either increasing or decreasing FR as demonstrated by a main effect of Stimulus Onset (Figure 3C). Interestingly, of the cells exhibiting interactions, 16 (14%) interacted with Context, whereas only 7 (6%) interacted with Pair. Figure 3D shows an example of a cell that increased firing when objects appeared but only when the context was the grey floor. Thus, a surprising number of POR neurons responded to object stimuli onset and some of those neurons showed conjunctive coding of stimulus onset with Context and/or Object Pair.

**Table 2.** POR neurons signal stimuli onset.

| Main effects and interactions by cell | Number | Percent |
| --- | --- | --- |
| StimOn | 25 | 21.9 |
| StimOn*Context | 12 | 10.5 |
| StimOn*Pair | 3 | 2.6 |
| StimOn*Context*Pair | 3 | 2.6 |
| StimOn*Context & StimOn*Pair | 1 | 0.9 |
| Signaled Onset | 44 | 38.6 |
| Did not Signal Onset | 70 | 61.4 |
| TOTAL | 114 | 100 |

Given that the context changed well before the object pairs were presented and that a choice had not yet been made during the analyzed epoch, we expected to see more conjunctive coding including Context compared with conjunctive coding including object Pair. To address this question, we conducted a frequency analysis based on the total numbers of main effects (Table S2B). Of the 44 cells that showed main effects and interactions with stimulus onset, there were 57 total main effects and interactions. The distribution of main effects and interactions were different for context and object pair. There were 16 (28.1%) significant interactions with context (StimOn*Context or StimOn*Context*Pair), whereas there were only seven (12.3%) interactions with Object Pair (StimOn*Pair or StimOn*Context*Pair). The number of interactions with Context was significantly greater than the number of interactions with object Pair ((X^2^(1) = 4.62, *p* < 0.032). The fact that context modulates responses to stimulus onset, even when the context changes earlier in time, is consistent with context operating as an occasion setter.

### POR neurons signal objects in particular locations and objects in particular contexts

In addition to the onset of floor contexts and the onset of object stimuli, we were interested in neuronal responses during task relevant behavioral epochs. We were especially interested in correlates of Context and Location given that in the nscBCD task context was predictive of correct choices, but location was not. For these analyses, we considered the location of the object chosen by the animal whether or not the choice was correct. We separately analyzed selectivity in the *stimulus*, *pre-choice*, and *post-choice* epochs (Figure 1D). For each cell and each epoch, we conducted a 2 X 4 X 4 analysis of variance (ANOVA) with Context, Location, and Object as the between-trial factors and firing rate as the dependent variable. For example, one condition might be the striped floor as Context and the animal‘s choice of Object 1, which had been presented in the north-east Location.

Selectivity was demonstrated by main effects and interactions among ‘Context’, ‘Location’, and ‘Object’. Again, we considered main effects to represent simple, non-conjunctive coding or selectivity and interactions to represent conjunctive coding or selectivity. When cells had an overlapping main effect and interaction, we presented only the interaction in Table 3A. For example, if a cell had a main effect of Context and a Context by Object interaction, this cell was counted under Interactions and was considered to exhibit conjunctive coding. However, if a cell had non-overlapping main effects and interactions, it would have been noted under Multiple Effects and Interactions in Table 3A, e.g. a cell with a main effect of Location and a Context by Object interaction would be considered to exhibit both conjunctive and non-conjunctive selectivity.

**Table 3A.** POR neurons show selectivity in three behavioral epochs.

| Significant Main Effects and Interactions | Stimulus Num(%) | PreChoice Num(%) | PostChoice Num(%) |
| --- | --- | --- | --- |
| MAIN EFFECTS |  |  |  |
| Context | 4(4) | 1(1) | 4(4) |
| Location | 22(21) | 36(39) | 27(28) |
| Object | 1(1) | 0(0) | 1(1) |
| INTERACTIONS |  |  |  |
| Object*Context | 9(9) | 0(0) | 5(5) |
| Object*Location | 5(5) | 4(4) | 2(2) |
| Context*Location | 6(6) | 8(9) | 5(5) |
| Context*Object*Location | 8(8) | 10(11) | 14(14) |
| MULTIPLE EFFECTS and INTERACTIONS |  |  |  |
| Context & Location | 3(3) | 1(1) | 5(5) |
| Location & Object | 2(2) | 0(0) | 1(1) |
| Context & Location & Object | 0(0) | 1(1) | 0(0) |
| Context*Object & Location | 3(3) | 5(5) | 9(9) |
| Context*Location & Object | 0(0) | 0(0) | 2(2) |
| Context*Location & Context*Object | 0(0) | 1(1) | 0(0) |
| Context*Object & Location*Object | 0(0) | 2(2) | 1(1) |
| Context*Object & Location*Object & Context*Location | 0(0) | 0(0) | 1(1) |
| Total selective cells | 63(61) | 69(75) | 77(81) |
| Total non-selective cells | 41(39) | 24(25) | 20(19) |
| TOTAL | 104 | 93 | 97 |

The stimulus epoch was the 500 ms when the objects first became visible. PreChoice was the 500 ms prior to approach to the object to signal a choice, and PostChoice was the 500 ms after the choice.

Of the 120 cells we recorded and isolated, 111 cells met criteria in at least one epoch, and 86 met criteria in all three epochs. In the *stimulus* epoch, 104 cells met criteria and 63 (61%) displayed selectivity (Table 3A). In the *pre-choice* epoch, 93 cells met criteria and 70 (75%) displayed selectivity. In the *post-choice* epoch, 97 cells met criteria and 79 (81%) displayed selectivity. Of the 111 cells that met criteria, 102 (92%) displayed selectivity in at least one epoch. Most cells had a single main effect or interaction: 55, 59, and 58 cells for the *stimulus*, *pre-choice*, and *post-choice* epochs, respectively. Fewer cells had multiple main effects and/or interactions: 8, 11, and 21 cells for the *stimulus*, *pre-choice* and *post-choice* epochs, respectively.

A central question of the present study was the extent to which the POR represents nonspatial context and whether representations involving context differed from those involving location. In the nscBCD task, context determined which object was correct and would be rewarded, but location was not predictive of reward. Thus, we were particularly interested in the distributions of main effects (non-conjunctive coding) vs interactions (conjunctive coding) involving Location and Context (Table 3B). These differences were examined by multiway chi-square analyses (Table S3B) for each epoch in which the primary grouping variable was statistical effects (main effects or interactions) and the secondary grouping variable was correlate (location and context). The outcome variable was whether or not the main effect or interaction was significant or nonsignificant.

**Table 3B.** POR main effects and interactions with Context vs Location.

| Main Effects and Interactions | Stimulus Num(%) | PreChoice Num(%) | PostChoice Num(%) |
| --- | --- | --- | --- |
| MAIN EFFECTS | *** | *** | *** |
| Context | 7(5) | 3(2) | 9(5) |
| Location | 30(21) | 43(27) | 42(22) |
| INTERACTIONS |  |  | # |
| Context | 26(18) | 27(17) | 38(20) |
| Location | 19(13) | 23(14) | 26(13) |
| INTERACTIONS WITH OBJECT |  |  | # |
| Context | 20(14) | 18(11) | 30(16) |
| Location | 13(9) | 14(9) | 18(9) |
| TOTAL EFFECTS & INTERACTIONS | 146 | 160 | 194 |

Percentages are of the total main effects and interactions for each epoch. Symbols: #, $p < 0.10$ ; \* $p < 0.05$ ; \*\* $p < 0.01$ ; \*\*\* $p < 0.001$ .

Even though the location of objects was not predictive of reward in this task, the location in which objects appeared was robustly represented with cells showing non-conjunctive (main effects) as well as conjunctive coding (interactions). Cells showed more main effects of Location than interactions with Location (Table 3B). Interestingly, quite a few cells showed selectivity for Location in the stimulus epoch when the rat was physically located near the middle of the maze. We observed three types of location selectivity including allocentric (east, west, north, and/or south), egocentric (left/right), and single locations. For some cells, preference for a single location was sustained across epochs (Figure 4A). Other cells showed interactions with context (Figure 4E left and middle panels; Figure 4F), objects (Figure 4E right panel), and with both objects and contexts (Figure 4D).

**Figure 4.**
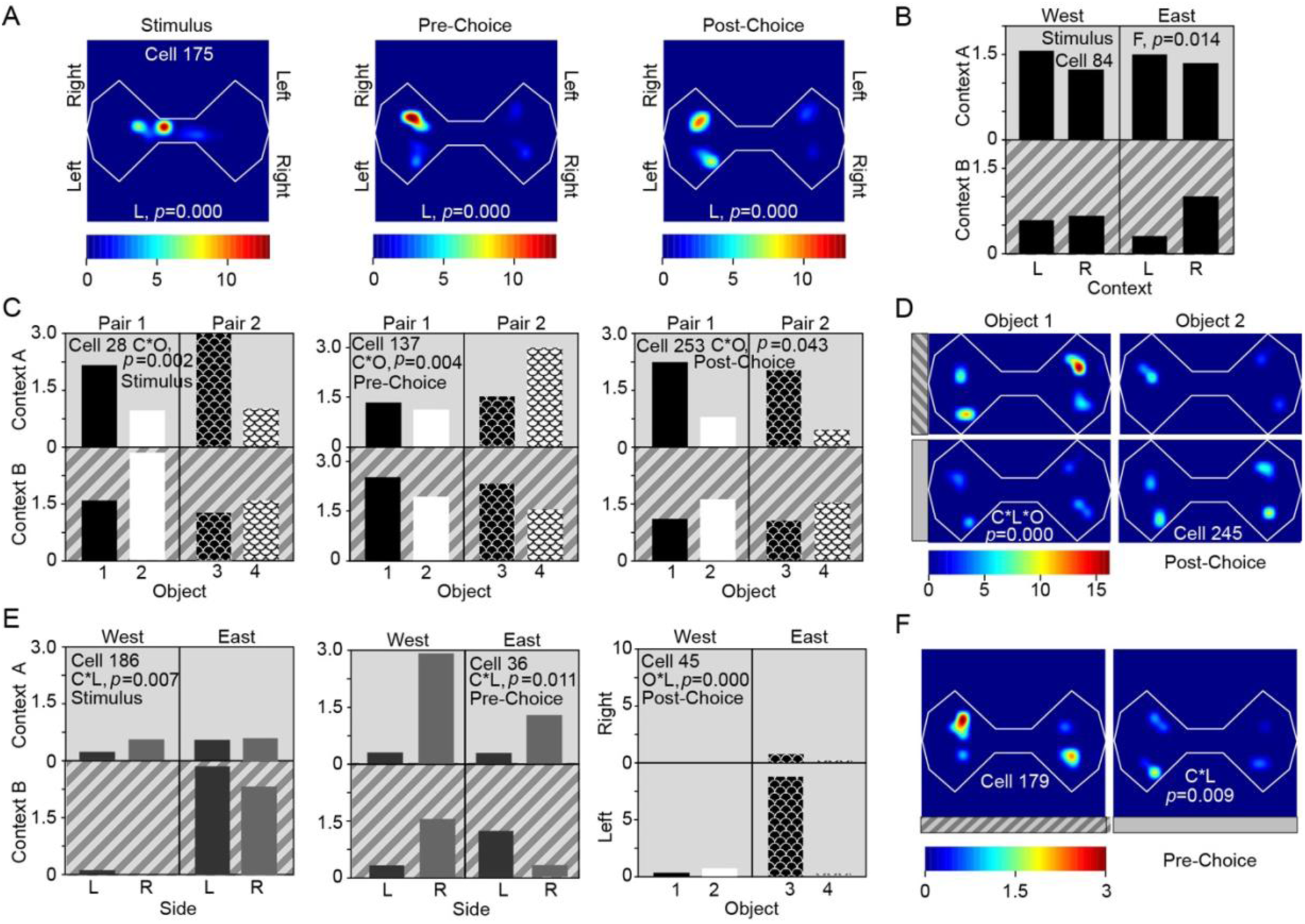
POR cells show conjunctive and non-conjunctive, task relevant coding for context, location, and object. Shown are representative cells in the POR that showed non-conjunctive main effects or conjunctive interactions among location (east-left, east-right, west-left, west-right), object (1,2,3,4), and context (A = striped, B = gray). Firing rates (FR) are denoted by heatmaps for spatial firing maps and are shown on the Y axis for histograms. (A) Spatial FR maps show an example of a cell with selectivity for a particular location sustained across the *stimulus*, *pre-choice,* and *post-choice* epochs. (B) Histogram for a cell showing non-conjunctive ‘Context’ selectivity as confirmed by a main effect of Context. Mean FR was higher for Context A (striped floor) regardless of location or stimuli. (C) Histograms for examples of ‘context x object’ (C*O) conjunctive coding. Cells in the left, center, and right histograms are from the *stimulus*, *pre-choice*, and *post-choice* epochs, respectively. For example, the cell in the left panel fired more for objects 1 and 3 in Context A and more for object 2 in Context B. (D) Spatial map for a cell with ‘context x location x object’ (C*L*O) conjunction. This cell responded preferentially when object #1 was presented in Context A on the egocentric left. (E) Histograms for examples of ‘context x location’ (C*L) conjunctions (left and middle) and an object by location (O*L) conjunction (right). Cells in the left, center, and right histograms are from the *stimulus*, *pre-choice*, and *post-choice* epochs, respectively. (F) Spatial FR maps for a cell with a C*L response.

Given that Context was predictive of reward, we were also interested in whether or not Context showed more conjunctive than non-conjunctive coding and whether Location showed a different pattern. Context showed more interactions than main effects and Location was more likely to show main effects than interactions. These differences were examined by multiway chi-square analyses (Table S3B) in which the primary grouping variable was epoch, the secondary grouping variable was correlate (location and context), and the outcome variable was presence or absence of significant interaction with object. Context exhibited significantly more conjunctive (interactions) than non-conjunctive coding (main effects) overall (X^2^(1) = 5.278, p <0.022). This pattern was evident and showed either marginally significant or significant associations for all three epochs: *poststim* (X^2^ (1)=3.632, p=0.057), *prechoice* (X^2^ (1)=15.882, p=0.001), and *postchoice* (X^2^ (1)=4.502, p=0.034). Figure 4B shows an example cell with a main effect of Context, and Figures 4C, D, E, and F show examples of cells demonstrating a variety of interactions involving context.

Interestingly, more cells showed Context*Object interactions (14%, 11%, 16%) than Location*Object interactions (9%, 9%, 9%) for *poststim*, *prechoice*, and *postchoice* epochs, respectively (Table 3B). Multiway chi-square with Epoch as the primary variable indicated a significant association. However, the association was not significant within the three epochs (p=0.196, p=0.456, and p=0.064, respectively).

Figure 4C shows examples of Context* Object conjunctive coding. One Context*Object cell showed conjunctive coding in the stimulus epoch, firing preferentially to objects 1 and 3 when the floor was striped and to object 2 when the floor was solid (Figure 4C, *left panel*). Another Context*Object cell fired preferentially to objects 1 and 3 when the floor was striped and to object 2 and 4 when the floor was solid during the pre-choice epoch (Figure 4C, *left panel*). In the post-choice phase, another Context*Object cell fired preferentially to objects 1 and 3 when the floor was striped and to object 2 and 4 when the floor was solid (Figure 4C, *left panel*). This could be interpreted as a reward cell. However, these analyses are based on the chosen object and include correct and incorrect choices.

Some cells showed Context*Location*Object interactions. For example, one cell fired equally for objects #1 and #2 in any location when the floor was gray. However, when the floor was striped, it fired preferentially to object #1 but only when it appeared on the egocentric left side (Figure 4D, *right panel*). Cells with a three-way interaction were observed in the *stimulus* (8 out of 104), *pre-choice* (10 out of 93), and *post-choice* epochs (14 out of 97). Taken together, these data are consistent with the hypothesis that POR cells encode the spatial layout of objects and patterns in the local physical context, even when location of objects is not relevant to the task.

### POR ensembles code for nonspatial context and object locations

Finally, we examined whether the ensemble coded for task relevant information by conducting representational similarity analyses (McKenzie et al. 2014). In this case we assessed the extent to which patterns of firing rates were similar across different conditions defined by the context (floor pattern), the identity of the object chosen, and the location of the chosen object. Firing rates for each recorded neuron were z-scored using the mean and standard deviation from the firing rates recorded during the behavioral epochs for each trial in the session. For each neuron and each epoch, a mean normalized rate was calculated for each of the 32 trial types defined by the three conditions (Context (2), Object (4), and Location (4)) resulting in 32 population vectors per epoch. The Pearson’s correlation coefficient was then calculated for each pair of population vectors for each trial epoch yielding a similarity matrix. For each condition, the magnitude of the effect was measured by comparing the mean correlation within a condition with the mean correlation across a condition for that component (for example: comparing the mean correlation between vectors with the same context and the mean correlation between vectors with different contexts). To estimate significance, a bootstrapped dataset was created using shuffled epoch assignments repeated 10,000 times and the correlations described above were recomputed for each shuffled set.

We found that the population activity showed significant correlations, defined as the difference of the mean correlations across trials with the same condition and trials with different conditions is greater than the 95^th^ percentile of the distribution of differences of the bootstrapped datasets. Location was significant during all epochs (Figure 5A). In the Pre-Context and Post-Context period, while the correlations between same-location trials were significantly different from different-location trials, this appears to be due to a higher anti-correlation between the different-location trials.

**Figure 5.**
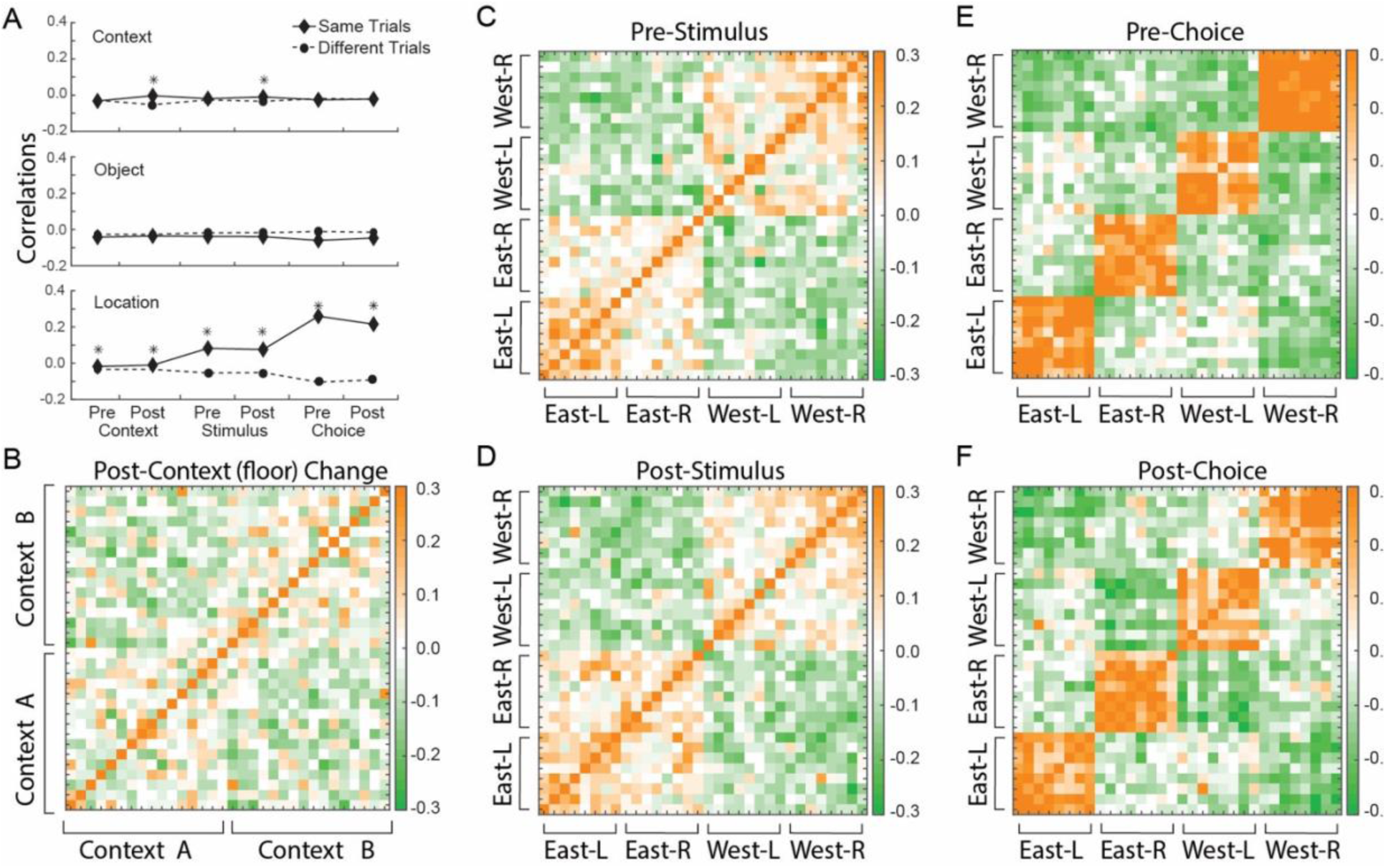
Postrhinal ensembles code for context and location in different epochs. A. Mean correlation coefficients for same-condition comparisons (solid line) and different-condition comparisons for the floor (top), object (middle), and object location (bottom). * indicates the difference between the solid and dotted line was greater than the 95^th^ percentile of the same measure from bootstrapped distributions. B. A correlation matrix using all cells across all sessions showing correlation coefficients by color code (right scale) for the 500 msec after the floor changed indicating a new context. The animal could be anywhere in the maze and the time to stimulus onset varies considerably. Context A and B are anticorrelated. C-D. Correlation matrices for pre-stimulus onset (C) and post-stimulus onset (D). The animal is in the center of the maze facing east or west. East and west are anticorrelated. E-F. Correlation matrices for pre-choice onset and post-choice onset. The animal is in one of the object location areas. The strongest anticorrelations are for combined opposite egocentric and allocentric locations. For example, West-Right is strongly anticorrelated with East-Left.

Using similarity matrices organized first by context (Figure 5B), we saw that during the Post-Context and Post-Stim periods there is correlation between similar contexts and anti-correlation between different contexts. This is interesting because the Context determines which object is correct, i.e., the Context is behaviorally relevant and becomes more salient when the objects appear. Looking at similarity matrices organized by object location (Figure 5C-F and Figure S1), we can see that during all epochs, allocentric locations were represented, and as the trial progresses representations of location become stronger. These representations appear unaffected by task relevant information. For example, there location differences for Pre- and Post-Context, for Pre- and Post-Stim, and for Pre- and Post-Choice. The broader allocentric locations of the east or west side of the maze were represented (Figure 5C, D), while the individual locations were represented more strongly during the pre- and *post-choice* periods (Figure 5E, F). Interestingly, during the *post-choice* period there was a strong anti-correlation with the location that was the furthest away in the combination allocentric and egocentric space, such as between the west-left location and the east-right location. Interestingly, Postrhinal ensembles show more evidence for conjunctive codes involving context than location in task relevant epochs (Figure S1).

## Discussion

We have proposed that the POR represents environmental contexts including the spatial layout of objects and patterns, that the POR automatically monitors the current context for changes and updates context representations when changes occur, and that object and pattern information arrives to the POR directly from the PER (Estela-Pro and Burwell, 2022, Furtak et al., 2012, Ho and Burwell, 2014, Peng and Burwell, 2021, Burwell and Hafeman, 2003). Such representations are available to multiple brain regions, for example, the hippocampus for associative learning (Agster and Burwell, 2013) and prefrontal cortex for context guided behavior and cognition (Hwang et al., 2018). We addressed two open questions about POR function. The first question was whether the POR represents nonspatial contexts by which we mean contexts that are not specifically tied to locations. The second question was whether POR representations of context modulate other cues in a manner consistent with occasion setting. To address these open questions, we recorded POR neurons as rats performed a nonspatial biconditional discrimination task in which the pattern on the floor, or the context, determined which of two objects was correct. The location of objects was not relevant to the discrimination. Thus, the experimental design allowed dissociation of location from non-spatial context (i.e. floor pattern).

### POR context representations operate like occasion setters

In our nscBCD task, we were able to compare neural correlates of context with neural correlates of location in three task relevant epochs – stimulus, pre-choice, and post-choice. We would expect POR neurons to exhibit both object-context conjunctive coding and object-location conjunctive coding. However, if POR representations of context operate as occasion setters, we would expect object-context conjunctive coding to be more frequent than object-location conjunctions because locations are not predictive of which object will be rewarded. For context, we found more conjunctive coding (interactions) than non-conjunctive coding (main effects), whereas for location, we found more non-conjunctive than conjunctive coding. Additionally, location was represented in more main effects than context was. Although location, is certainly a component of the context, in this task the floor pattern interacts with other cues in a way that is more consistent with how contexts have been suggested to work in that modulate the meanings of other cues in an environment (Fraser and Holland, 2019). The patterns of conjunctive and non-conjunctive coding we observed for the predictive context and the nonpredictive location are consistent with two interpretations. First POR does appear to represent the nonspatial context operationalized as a pattern on the floor, and the data are consistent with an occasion setting interpretation of how POR representations of context interact with other features of the environment.

Thus far we have examined our findings in the framework of occasion setting in which the occasion setter (floor pattern in this case), interacts with other cues in a modulatory process that is distinct from a simple associative process (Fraser and Holland, 2019). It might also be useful to examine whether our findings might differentiate between alternative theories of occasion setting might account for our findings.

### POR object-context coding is not related to learning

In prior work we established that the POR has a role in processing contextual information (Bucci et al., 2021, Bucci et al., 2002, Burwell et al., 2004a, Burwell et al., 2004b, Heimer-McGinn et al., 2017). We also reported that the POR exhibits object-location conjunctive coding (Furtak et al., 2012), which can be viewed as a signature of context representations. In that study, in which location was also not predictive, 10-15% of cells showed object-location coding depending on the epoch. In the present study, 13-20% showed object location conjunctive coding across the same three epochs. The number of cells with non-conjunctive location correlates was also similar with the Furtak et al. (2012) study reporting 11-44% depending on the epoch, and the present study reporting 29-46%. Thus, our findings regarding POR object location correlates when location does no predict outcome was largely replicated.

It is also useful to compare our findings with those of a study in which location was predictive of the correct choice. McKenzie et al. (2014) recorded hippocampal neurons in a similar context biconditional discrimination task, except that the two spatial contexts differed in allocentric location as well as other environmental cues. On the east side of the maze, one odor of a pair was correct and on the west side of the maze the other odor was correct. That study reported that 74.7% of isolated cells were influenced by allocentric location (40.7%, east vs west location of trial) or egocentric location of the samples (34.0%, left vs right location of odor). A total of 13.5% of cells coded for odor identity alone, and 42.7% showed conjunctive coding of odors with spatial context. In our study, taking into consideration all three task relevant epochs, 42.6% of isolated cells were influenced by nonspatial context (floor pattern), 78.7% were influenced by egocentric and/or allocentric location. 40.7% coded for object identity conjunctively or non-conjunctively, and 36.1% showed conjunctive coding of objects with context or location. START Thus, we found that a higher number of cells coded for object identity, but the majority was conjunctive coding, and only 9.3% of isolated cells showed non-conjunctive coding in the form of main effects of object identity, which may be more comparable to the McKenzie et al. (2014) study. We also observed a higher proportion of cells influenced by location. In our study, however, location cells included egocentric and allocentric correlates, whereas location correlates in the McKenzie and colleagues study included only the left vs right position of item stimuli. If we consider that perhaps half of our location correlates signaled egocentric locations, then the numbers are roughly comparable. Importantly, McKenzie et al. (2014) reported that 40.7 percent of recorded hippocampal cells were influenced by spatial context, whereas we found that 42.7% of recorded POR cells were influenced by non-spatial context.

One interesting question about object-context coding in the POR and item-location coding in the hippocampus is the timing of the emergence of such codes. Using the same task as McKenzie and colleagues, Komorwski et al. (2009) reported that item-location coding emerged as rats learned the task. In that study, the proportion of active cells with item-location correlates increased during learning from 6.4% in the first 30 trial block to 25% in the second block to 31.3% in the third 30 trial block. Over the same blocks, percent correct increased from about 45.5% to 79.4% to 84.8%. Thus, the proportions of item-location selective cells predicted accuracy. In our study, rats performed at 55-65% correct throughout recording and proportions of object-context cells were stable across sessions at about 32.6% of active cells and up to 42.4% if we include object-location cells. This suggests that object-context coding in the POR does not depend on learning and does not predict accuracy. These findings are consistent with the hypothesis that the POR is broadly and automatically representing both spatial and nonspatial contexts and making those representation available to other brain regions, including the hippocampus. The hippocampus, then, uses these representations to form context-guided object associations (Komorowski et al., 2009, McKenzie et al., 2014).

Interestingly, although location was not predictive of reward, the locations in which objects appeared were robustly represented by POR neurons. When animals were in the middle of the maze just after the onset of objects, east and west were strongly anticorrelated. In subsequent epochs, all four locations were represented. Interestingly the most different locations east-left vs west-right and east-right vs west-left in that they were the most anticorrelated in the ensemble analyses. Thus, the POR appears to be mapping task-relevant locations in the context.

### POR monitors the current context for changes

Our study also provides evidence that the POR monitors the current context for changes. We found that about half of POR neurons signaled when the floor changed, and about half of those cells signaled the appearance of a particular floor. We also found that POR neurons signaled the presentation or onset of the object pairs to be discriminated. Notably, when the objects appeared, cells were more likely to show conjunctive coding involving context as opposed to conjunctive coding involving the identity of the object pairs. This is surprising given that the context would have changed well before the objects were presented and that object presentation was a highly salient event. These findings are consistent with the idea that the POR monitors the current context for changes and updates the representation of context when changes occur.

### Spatial and non-spatial components of context

A growing body of evidence indicates that visual context includes both spatial and non-spatial elements, and that the POR/PHC is involved in processing both types of information (Mullally and Maguire, 2011). In rodent and non-human primate work, for instance, it is common to examine context as defined by background images (Bachevalier et al., 2015), patterns on the floor (Heimer-McGinn et al., 2017), or patterns on the wall (Norman and Eacott, 2005, Park et al., 2017). In these studies, lesions to the POR impair context-guided behaviors in the absence of location cues. Beyond simple visual cues, human studies have defined context as a movement contingency (stable vs in motion) (Karanian and Slotnick, 2014),and as a specific written question (question defines trial type) in a lexical task (Diana, 2017). In these studies, PHC is preferentially activated during contextual processing. Other studies go even further, showing that POR/PHC is involved in contextual processing as defined by motivational state (hungry vs satiated) (Burgess et al., 2016, LaBar et al., 2001). Taken together, these studies provide strong evidence that processing of context in POR/PHC goes beyond location and spatial layout. Our study is the first to investigate the neural correlates of this non-spatial contextual processing, suggesting that POR neurons bind together streams of spatial and non-spatial contextual information.

### Cortical integration of object and spatial information

For some years, now, there has been a focus on segregated spatial and nonspatial processing streams that provide information to the hippocampus. More specifically, the PER provides nonspatial input to the lateral entorhinal cortex and the POR provides spatial input to the medial entorhinal cortex. In fact, available data argues against such a segregation. For example, there is considerable evidence that the PER processes spatial and contextual information (e.g. Bucci et al., 2021, Heimer-McGinn et al., 2017, Lim et al., 2022). Anatomically, we have known for some time that, although the PER is preferentially connected with the lateral entorhinal cortex. Often overlooked, the POR is robustly interconnected with *both* the lateral and the medial entorhinal cortices (Burwell and Amaral, 1998, Doan et al., 2019). Moreover, the rodent POR and the primate PHC are robustly and reciprocally connected with the PER (Burwell and Amaral, 1998, Suzuki and Amaral, 1994), which could explain the presence of object information in the POR.

In addition to the anatomy, other lines of evidence indicate that object information is processed in the POR. Although the POR/PHC is not necessary for object discrimination (Norman and Eacott, 2005), object-in-place and object-in-context discriminations do rely on POR/PHC, as indicated by lesion studies in rodents and primates (Norman and Eacott, 2005, Park et al., 2017, Malkova and Mishkin, 2003, Bachevalier et al., 2015). These findings are also supported by human imaging studies showing PHC activation during similar discriminations (Karanian and Slotnick, 2017, Karanian and Slotnick, 2014, Yeung et al., 2019, Faivre et al., 2019, Stevenson et al., 2020). Importantly, disconnection of the POR and PER impairs object-in-context discrimination, suggesting that POR may require object input from the PER in order to perform this task (Heimer-McGinn et al., 2017). Moreover, molecular studies reveal arc activation in POR during both spatial and non-spatial tasks (Beer et al., 2013), and demonstrate that delivery of protein kinase C to POR enhances shape discrimination (Zhang et al., 2019, Zhang et al., 2017). Finally, a handful of studies report directly on the neural correlates of object processing in POR. These studies discuss the presence of POR correlates for object-location conjunctions (Furtak et al., 2012) and for object movement (Nishio et al., 2018, Beltramo and Scanziani, 2019). Taken together, these studies support the idea that information about objects is bound to context in the POR.

### POR monitors the layout of scenes and contexts

Our results indicate that cells in the POR respond to changes in the visual layout of a scene and that this response may depend on salience of the change. An appropriate analogy might be a familiar empty room with gray walls. If patterned wallpaper was added, or if two large pieces of furniture were added, the room would feel quite different. Our data demonstrates that POR cells responded to these types of highly salient changes, specifically, changes to floor pattern and object onset. In addition, 25% of cells were tuned to floor changes by floor identity, meaning they responded only to gray-to-stripe changes or to stripe-to-gray. This makes sense considering that rats were highly familiar with both contexts. In fact, there is some data to suggest that POR representations may be stable within testing sessions (Furtak et al., 2012) and possibly as long as 2-10 days (Burgess et al., 2016). This may account for the ability of POR cells to discriminate the *direction* of the floor change between two highly familiar contexts. POR cells also discriminated between more subtle changes, such as object onset in a specific context (10%; new furniture in gray room vs newly painted room), onset of particular objects (3%; new red vs blue armchairs of similar size/shape), and onset of particular objects in a specific context (3%; red armchairs in the gray room only). In each of these analogous situations, the differences between layouts are more subtle than gray vs wallpaper (floor change) or empty vs furnished (object onset). It makes sense, then, that less cells would respond to these less pronounced changes. It very well could be that the number of responsive cells corresponded to how salient the change or difference between layouts was. Thus, our results support the hypothesis that the POR monitors the current layout of a scene, including objects and non-spatial context, and that it is responsible for signaling when the layout changes.

## Conclusion

Our study shows that cells in the POR process non-spatial context and that POR context representations can be used to process the meaning of other available cues. The nonspatial context, in this case, was the pattern on the floor that predicted which of two objects would be rewarded. Although location was part of our nscBCD task in that objects did appear in particular locations, location was not predictive of reward. We observed significantly more object-context conjunctive coding compared with object-location conjunctive coding. These observations are consistent with the idea that context representations operate as occasion setters by setting the meaning of other cues. Object-context conjunctive coding was not predictive of learning or accuracy as has been observed in the hippocampus. We also observed that the POR signaled the onset of context changes and the onset of objects. This finding is consistent with the idea that the POR is monitoring the current environment for changes and updating the current representation of context. This study provides further evidence that the object information is processed by the POR, but primarily in the service of representing the spatial layout of objects and patterns that characterize the current context. Taken together, the available evidence suggests that 1) the POR is important for representing spatial context including the geometry of the space and the spatial layout of objects and patterns, 2) the POR codes for contexts that are distinguished only by non-spatial features e.g. surface patterns, 3) object and pattern information necessary for both spatial and nonspatial representations likely arrive to the POR directly from PER, and 3) the POR signals changes to the current context. Future work should address the mechanisms by which the POR represents context and monitors changes.

## Detailed Methods

### Contact for reagent and resource sharing

Further information and requests for resources and reagents should be directed to and will be fulfilled by the Lead Contact, Rebecca D. Burwell.

### Experimental model and subject details

Subjects were five adult Long Evans rats (Charles River Laboratories, Wilmington, MA), singly housed in diurnal conditions (12 h light/dark cycle) with *ad libitum* access to food and water. Before surgery rats were handled at least three times per week and maintained at 85– 90% body weight. At the time of surgery, all subjects were 3–5 months old and weighed 250– 300 g, and had no involvement in previous procedures. After surgery, rats were allowed to recover and then handled daily. Rats were group housed before surgery (2-3 per cage) and single housed after surgery to prevent damage to the implanted devices. Supervised play time with previous cage mates was allowed for at least 1 hr per day. All procedures were in accordance with NIH guidelines for the care and use of rats in research. The protocol covering these experiments was approved by the Brown University Institutional Animal Care and Use Committee.

### Method details

#### Apparatus

Testing was performed in a bowtie-shaped arena (147 x 111 cm) made of white matte acrylic and placed over a Floor Projection Maze (Jacobson et al., 2014, Furtak et al., 2009, Furtak et al., 2012), in which images are back-projected to the floor. The apparatus was housed in an isolated testing room monitored by an overhead video camera and interfaced with integrated systems for neuronal data acquisition, location tracking, and behavioral control. Reward was delivered by inter-cranial stimulation (ICS) in the form of bipolar electrical pulses to the medial forebrain bundle. ICS pulses were 500 μsec in length, separated by 500 μsec, and sustained for 500ms. Pulse frequency was fixed at 100 Hz. Amplitude ranged 15-35 μA depending on the sensitivity of the subject. The stimuli were pairs of 2D objects measuring 12.5 cm in diameter spaced 24 cm apart and presented on the floor, which is ideal considering the visual acuity limits for Long Evans rats (Furtak et al., 2009). All pairs of objects were luminance-matched to ensure that shape was the salient cue. Context was defined visually by floor pattern and was different on each trial; floor A was striped grey and floor B was solid gray. The behavioral program, including white noise, floor image displays, and ICS delivery, were controlled by DIG-716P2 Smart Control Output Interface (Med Associates Inc, St. Albans, VT) using a custom-written script in Med-PC IV software. Trial information stored in Med-PC output files was extracted using custom MATLAB scripts (Mathworks).

#### Stimuli

The visual stimuli consisted of two floor patterns and three pairs of objects. All pairs of objects were luminance-matched to ensure that shape was the salient cue. Existing literature on rats’ visual acuity indicates that pigmented rats can discriminate approximately 1 cycle per degree (cpd), or 30 arc minutes (Lashley, 1930, Prusky et al., 2002). The stimuli were 12.5 cm in diameter and were presented at a distance of 24 cm, occupying >1700’ of the visual field. The smallest feature of the shapes was the dark borderof the shapes, which is 0.42 cm at the proscribed viewing distance of 24 cm. The angular size corresponds to 51 arcminutes, well within the 30 arc-minute threshold.

Previous literature and data from our lab indicate that pigmented Long-Evans rats can detect differences in grey with contrast differences as low as 12% on a linear brightness scale such that white is 0% and black is 100% (Furtak et al., 2009, Powers and Green, 1978). To ensure that subjects in this task were able to distinguish between the two different floor patterns, the stripes of the striped floor were designed to be 27% grey and 61% grey, whereas the solid floor was designed to be 44% grey. To provide the greatest possible level of contrast, the interior of each of the 2D objects was white (0%) with outlines that were 91% grey.

#### Surgery

Rats were anesthetized with isoflurane and implanted with custom-designed microdrives built in-house and containing up to 32 individually drivable stereotrodes (25 μm nichrome wires, A-M Systems, Inc., Carlsborg, WA). The incisor bar was adjusted such that the bregma and lambda were in the same horizontal plane (±0.2 mm). A 2 mm craniotomy was made in the left or right hemisphere at −0.5 mm anterior and 5.0 mm lateral to lambda, allowing for visualization of the sagittal sinus. The electrodes were inserted at a 16° angle along the ML axis with tip pointed in the lateral direction and lowered 3.5 mm relative to the cortical surface just anterior to the sinus. Two ICS electrodes were implanted −2.5 mm posterior and ±2.0 mm lateral to bregma and lowered 8.5 mm relative to skull. Dental cement and anchor screws were used to secure the electrode assembly to the dorsal surface of the skull. Following surgery, rats were allowed 3 days to recover and then each stereotrode was advanced daily in increments of 1/8 turn (∼20 μm).

#### Behavioral Training

In this study we present a contextually guided bi-conditional discrimination task (nscBCD), in which location and context can be dissociated. Two pairs of 2D objects, pair 1 and pair 2, were randomly intermixed in every session. In each trial, the rats were presented with one of two pairs of objects on one of two different floor patterns in a bowtie shaped maze. The floor patterns determined which object was correct within each pair. The pairs of objects were presented on either side of the bowtie maze, and the animals were rewarded for approaching the correct object. The location of the objects was counterbalanced in both egocentric (left vs right) and allocentric (east vs west) space. Thus, correct choices in this task required associating the floor pattern with the rewarded object, but location did not modify the likelihood of *post-choice*. An incorrect choice was followed by a correction trial with the same parameters (e.g. pair 1, floor A, correct on left) on the opposite side of the bowtie maze. The number of consecutive correction trials was limited to prevent spatial biases.

Before surgery, rats were habituated to the room and apparatus. After surgery, rats were shaped to ICS reward, to stop in “ready” position, and to approach the stimulus. After shaping, rats were sequentially trained on simple and context discrimination (using training objects), and finally on pairs 1 and 2. For each stage of training, rats were expected to meet trials to criterion (TTC) for two days in a row before moving on to the next stage. TTC was defined as 10 ten correct choices in 12 consecutive trials. In all phases of shaping and training in which two items were presented, the left versus right location of the correct stimulus was counterbalanced. At each stage, the number of trials per session was increased gradually until the animals were able to complete 200 trials per session. Once training was complete, rats received one daily session of 200 trials with pairs 1 & 2 and floors A & B.

#### Electrophysiology

Neuronal signals were recorded from stereotrodes, amplified with a gain of 20 at the headstage (HST/32V-G20, Plexon, Inc., Dallas, TX), and passed through a differential pre-amplifier with a gain of 50 (PBX2, Plexon, Inc.). Single unit activity was filtered between 154-8,800 Hz and digitized at 40 kHz. LFPs were filtered between 0.7–170 Hz and digitized at 1 kHz. Signals were further amplified for a total gain of 10,000 (MAP system, Plexon, Inc.). Two rats were recorded using the Plexon OmniPlex® system. Signals were collected through a 64-channel digital headstage (HST/64D, Plexon, Inc., Dallas, TX). Single units were filtered between 0.77–6000 Hz and up sampled at 40 kHz through a digital headstage processor (DPH). LFPs were filtered at 0.7-170 Hz and digitized at 1 kHz. For all rats, waveforms were extracted by real-time thresholding (Sort Client, Plexon, Inc.).

#### Histology

After recording was concluded, animals were given an overdose of Beuthanasia-D (100 mg/kg, intraperitoneal) and a small marking lesion (1-15 μA current for 15 sec) was made at the end of each stereotrode. Rats were pericardially perfused with 0.1M phosphate-buffered saline (PBS) followed by 10% formalin. Brains were cryoprotected in 30% sucrose for 3-5 days prior to sectioning. Brains were cut in the coronal plane (60 μm), mounted on charged slides, stained with thionine, and cover slipped with DPX. The locations of electrode tips were reconstructed with a light microscope.

### Quantification and statistical analysis

#### Unit Isolation

Recorded units were sorted using a variety of manual and partially automated techniques for isolation based on waveform characteristics in Offline Sorter (Plexon, Inc). Only cells recorded during sessions with at least 55% accuracy on the behavioral task were considered. Furthermore, only well-isolated units with a signal-to-noise ratio greater than 2:1, and spike separation of at least 1 ms were retained for further analysis. Overall, we isolated 120 cells from 5 animals across 54 sessions. Time alignment was corrected based on a known issue between spike data and field potential data using FPAlign (Plexon, Inc) (Nelson et al., 2008). Spike timestamps were extracted from sorted files using NeuroExplorer and used to compute FR across key behavioral events (NEX; Plexon, Inc).

#### Factorial Analyses of Neural Correlates

In order to assess the neuronal correlates of behaviorally relevant event markers we ran a between-trials factorial ANOVA for each cell for three epochs (Table 3A; Figure 4). For each cell, neuronal activity (spikes/sec) was analyzed for three epochs: *stimulus* (500 ms after onset of object stimuli), *pre-choice*, and *post-choice* (500 ms before and after rat makes a choice). Between-trial variables were ‘object pair’ (pair 1 or pair 2), ‘Context’ pattern (stripe or solid), ‘Location’ (east-left, east-right, west-left, and west-right). Analyses were performed in SPSS Version 24. Main effects and interactions were considered significant at *p*-value < 0.05, and cells were included for analysis if they fired at least 30 times within a given epoch.

#### Floor Change Analysis

The floor change analysis was conducted to examine the effects of floor change on FR (Figure 3). For each cell, we compared the FR 500 ms before (*prefloor)* to 500 ms after (*postfloor)* a change in floor. A repeated measures ANOVA was carried out with ‘epoch’ as the within-trial factor (*prefloor* vs *postfloor*) and ‘Context’ as the between-subject factor (striped or gray). Analyses were performed in SPSS Version 24 and main effects and interactions were considered significant at a *p*-value < 0.05. Cells were included if they fired at least 20 times within a given epoch.

#### Stimulus Onset Analysis

The object onset analysis examined whether FR changed in response to onset of a 2D object. Similarly to the floor change analysis, FRs were compared across epochs that occurred 500 ms before (*prestim)* and after (*poststim)* stimulus onset. For each cell, a repeated measures ANOVA was carried out with ‘epoch’ as the within-trial variable (*prestim* vs *poststim*), and ‘Context’ (gray or striped) and ‘pair’ (object pair 1 or 2) as the between-trial variables. Analyses were performed in SPSS Version 24 and main effects and interactions were considered significant at a *p*-value < 0.05. Cells were included if they fired at least 20 times within a given epoch. Out of the 120 isolated cells, 114 met analysis criterion for this analysis.

#### Similarity Analyses

The similarity matrices were calculated in the same manner as in (McKenzie et al. 2014). In this case we assessed the extent to which patterns of firing rates were similar across different conditions within three components: context (floor pattern), the identity of the object chosen, and the location of the chosen object. Firing rates for each recorded neuron were z-scored using the mean and standard deviation from the firing rates recorded during each of the six behavioral epochs for each trial in the session. The 500 ms epochs included Pre- and Post-Context (floor pattern) onset, Pre- and Post-Object onset, and Pre- and Post-Choice of an object. For each neuron and each epoch, a mean normalized firing rate was calculated for each of the 32 trial types defined by components and conditions: (Context (2), Object (4), and Location (4)). We then constructed population vectors for each of the 32 component conditions for each of the six epochs. For example, in the pre-stimulus epoch one of the 32 population vectors would comprise the mean firing rate for each neuron in the ensemble for trials in which the component conditions were Context A, Object 1, and Location 1. The Pearson’s correlation coefficient was then calculated for each pair of population vectors yielding a 32 x 32 similarity matrix for each trial epoch.

For each condition, the magnitude of the effect was measured by subtracting the mean correlation across conditions from the mean correlation within a condition with for a particular component. For example, in the case of Context, the mean correlation between vectors with the different contexts (A vs A and B vs B) was subtracted from the mean correlation between vectors with same contexts. To estimate significance, a bootstrapped dataset was created using shuffled epoch assignments repeated 10,000 times and the correlations described above were recomputed for each shuffled set. The population firing rate was said to be significantly correlated with a component (Context, Location, or Object) when the magnitude of the effect was greater than the 95^th^ percentile of the distribution of effect magnitudes from the bootstrapped datasets.

## Acknowledgements

This work was funded by NIMH R01MH108729, NSF IOS 1656488, and NSF IOS-1146334 to RDB; NIMH F32 MH105210 to VHM; and NEI F32 EY031561 to SGT. VHM is now located at Department of Psychology, Roger Williams University, Bristol, RI 02809.

## Author Contributions

Victoria R. Heimer-McGinn: investigation, software, formal analysis, data curation, visualization, writing - original draft as well as review and editing. Sean G. Trettel: software, data curation, formal analysis, writing - review and editing. Brendon Kent: Investigation, software, writing - review and editing. Amrita N. Singh: investigation, visualization, writing - review and editing. Rebecca D. Burwell: conceptualization, supervision, project administration, funding acquisition, methodology, software, formal analysis, writing - review and editing, visualization, supervision.

## Supplemental Materials

**Table S1:**
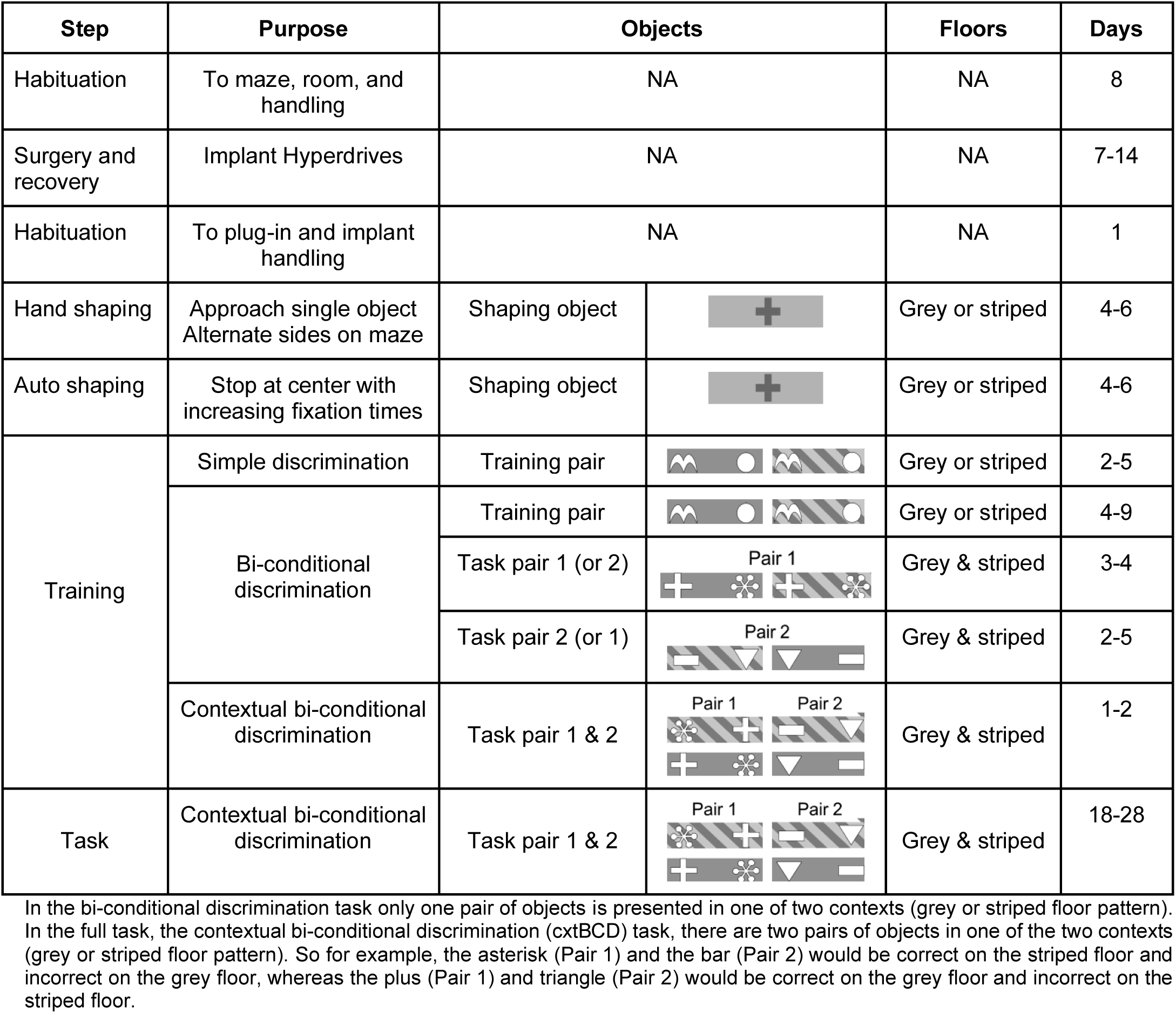
Habituation and Shaping Steps.

| Step | Purpose | Objects |  | Floors | Days |
| --- | --- | --- | --- | --- | --- |
| Habituation | To maze, room, and handling | NA |  | NA | 8 |
| Surgery and recovery | Implant Hyperdrives | NA |  | NA | 7-14 |
| Habituation | To plug-in and implant handling | NA |  | NA | 1 |
| Hand shaping | Approach single object<br>Alternate sides on maze | Shaping object |  | Grey or striped | 4-6 |
| Auto shaping | Stop at center with increasing fixation times | Shaping object |  | Grey or striped | 4-6 |
| Training | Simple discrimination | Training pair |  | Grey or striped | 2-5 |
|  | Bi-conditional discrimination | Training pair |  | Grey or striped | 4-9 |
|  |  | Task pair 1 (or 2) | Pair 1 | Grey & striped | 3-4 |
|  |  | Task pair 2 (or 1) | Pair 2 | Grey & striped | 2-5 |
|  | Contextual bi-conditional discrimination | Task pair 1 & 2 | Pair 1 Pair 2 | Grey & striped | 1-2 |
| Task | Contextual bi-conditional discrimination | Task pair 1 & 2 | Pair 1 Pair 2 | Grey & striped | 18-28 |

In the bi-conditional discrimination task only one pair of objects is presented in one of two contexts (grey or striped floor pattern). In the full task, the contextual bi-conditional discrimination (cxtBCD) task, there are two pairs of objects in one of the two contexts (grey or striped floor pattern). So for example, the asterisk (Pair 1) and the bar (Pair 2) would be correct on the striped floor and incorrect on the grey floor, whereas the plus (Pair 1) and triangle (Pair 2) would be correct on the grey floor and incorrect on the striped floor.

**Table S2A.**
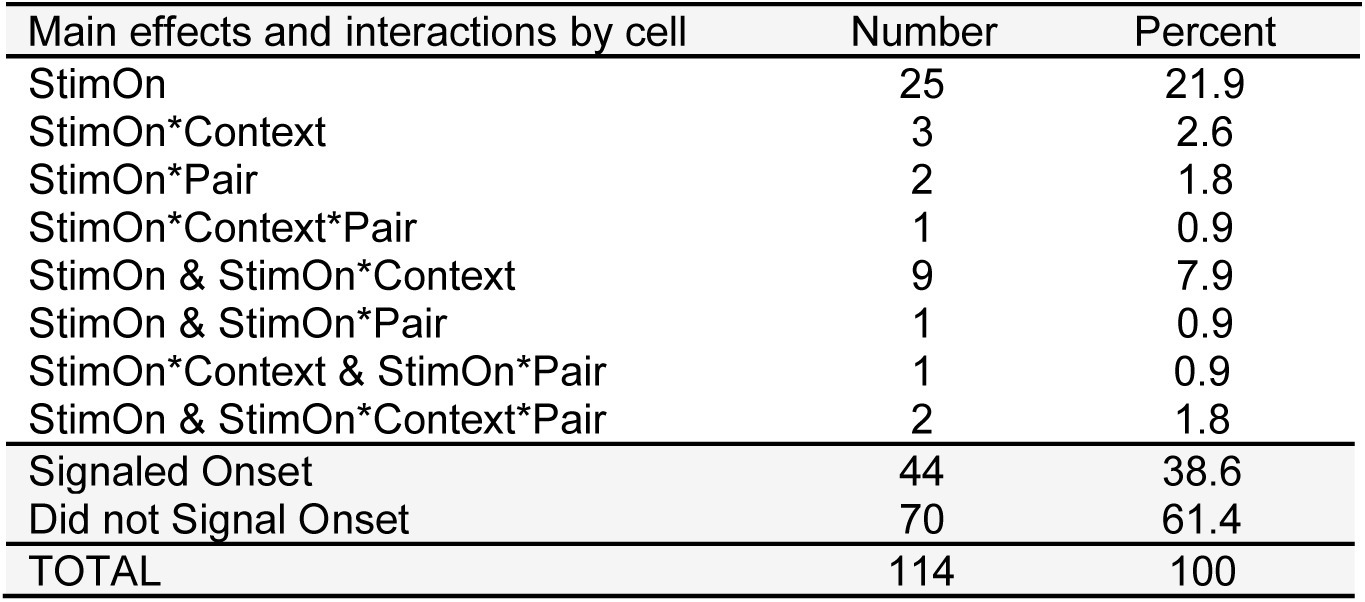
POR neurons signal stimuli onset.

| Main effects and interactions by cell | Number | Percent |
| --- | --- | --- |
| StimOn | 25 | 21.9 |
| StimOn*Context | 3 | 2.6 |
| StimOn*Pair | 2 | 1.8 |
| StimOn*Context*Pair | 1 | 0.9 |
| StimOn & StimOn*Context | 9 | 7.9 |
| StimOn & StimOn*Pair | 1 | 0.9 |
| StimOn*Context & StimOn*Pair | 1 | 0.9 |
| StimOn & StimOn*Context*Pair | 2 | 1.8 |
| Signaled Onset | 44 | 38.6 |
| Did not Signal Onset | 70 | 61.4 |
| TOTAL | 114 | 100 |

**Table S2B.** POR cells show more conjunctive coding with context than with object pair.

| Total Main Effects and Interactions over 44 selective cells | Number of Effects | Percent |
| --- | --- | --- |
| StimOn | 37 | 64.9% |
| StimOn*Context | 13 | 22.8% |
| StimOn*Pair | 4 | 7.0% |
| StimOn*Context*Pair | 3 | 5.3% |
| TOTAL EFFECTS | 57 | 100.0% |
| Interactions with Context | 16 | 28.1% |
| Interactions with Pair | 7 | 12.3% |

Chi Square: Stimulus Onset Interactions with Context vs Pair.

Chi Square: Stimulus Onset Interactions with Context vs Pair
|  | Interactions | No Int. | Statistics |  |
| --- | --- | --- | --- | --- |
| Context | 16 | 41 | Chi Square | 4.4118 |
| Pair | 7 | 50 | p-value | 0.03569 |

**Table S3B:**
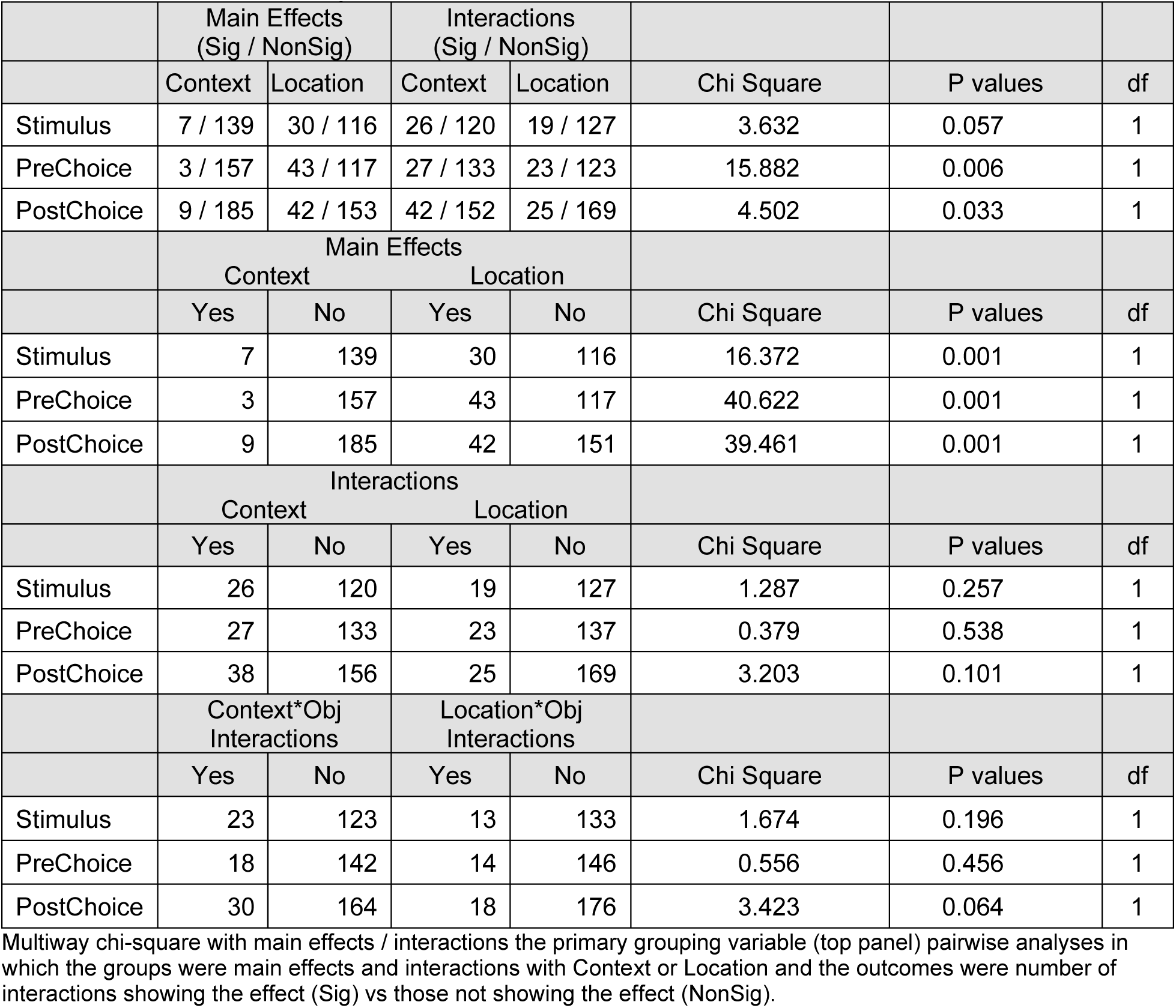
Statistical analyses for Table 3B.

**Figure S1.**
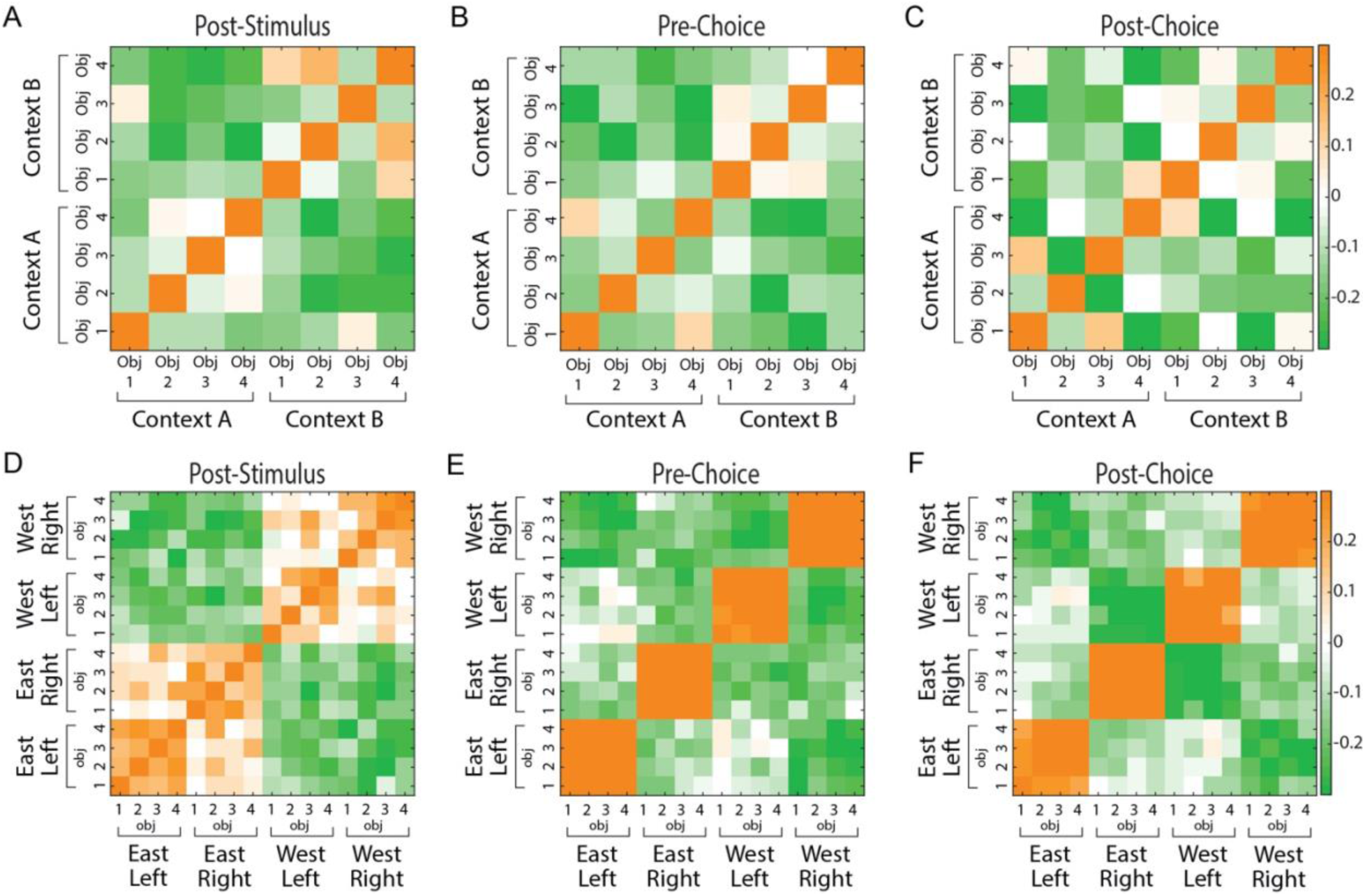
Postrhinal ensembles show more evidence for conjunctive codes involving context than location in task relevant epochs. Shown are correlation matrices using all cells across all sessions showing correlation coefficients by color code (right scale) for the 500 msec of three task relevant epochs: post-stimulus (A, D), pre-choice (B, E), and post-choice (C, F). The upper panels (A, B, C) show matrices for objects within context. Although most cells are anticorrelated, especially across contexts, there are some correlates across objects within a context. The lower panels (D, E, F) show matrices for objects within locations. Here, objects do not appear to interact with location at all.

## References

Agster, K. L. & Burwell, R. D. 2013. Hippocampal and subicular efferents and afferents of the perirhinal, postrhinal, and entorhinal cortices of the rat. Behav Brain Res, 254, 50–64.

Aguirre, G. K. & D’esposito, M. 1999. Topographical disorientation: a synthesis and taxonomy. Brain, 122 (Pt 9), 1613–28.

Aminoff, E. M., Kveraga, K. & Bar, M. 2013. The role of the parahippocampal cortex in cognition. Trends Cogn Sci, 17, 379–90.

Bachevalier, J., Nemanic, S. & Alvarado, M. C. 2015. The influence of context on recognition memory in monkeys: effects of hippocampal, parahippocampal and perirhinal lesions. Behav Brain Res, 285, 89–98.

Bar, M. & Aminoff, E. 2003. Cortical analysis of visual context. Neuron, 38, 347–58.

Beer, Z., Chwiesko, C., Kitsukawa, T. & Sauvage, M. M. 2013. Spatial and stimulus-type tuning in the LEC, MEC, POR, PRC, CA1, and CA3 during spontaneous item recognition memory. Hippocampus, 23, 1425–38.

Beltramo, R. & Scanziani, M. 2019. A collicular visual cortex: Neocortical space for an ancient midbrain visual structure. Science, 363, 64–69.

Bonner, M. F. & Epstein, R. A. 2021. Object representations in the human brain reflect the co-occurrence statistics of vision and language. Nat Commun, 12, 4081.

Bucci, D. J., Phillips, R. G. & Burwell, R. D. 2000. Contributions of postrhinal and perirhinal cortex to contextual information processing. Behav Neurosci, 114, 882–94.

Bucci, D. J., Phillips, R. G. & Burwell, R. D. 2021. Contributions of postrhinal and perirhinal cortex to contextual information processing. Behav Neurosci, 135, 313–325.

Bucci, D. J., Saddoris, M. P. & Burwell, R. D. 2002. Contextual fear discrimination is impaired by damage to the postrhinal or perirhinal cortex. Behav Neurosci, 116, 479–88.

Burgess, C. R., Ramesh, R. N., Sugden, A. U., Levandowski, K. M., Minnig, M. A., Fenselau, H., Lowell, B. B. & Andermann, M. L. 2016. Hunger-Dependent Enhancement of Food Cue Responses in Mouse Postrhinal Cortex and Lateral Amygdala. Neuron, 91, 1154–1169.

Burwell, R. D. & Amaral, D. G. 1998. Perirhinal and postrhinal cortices of the rat: interconnectivity and connections with the entorhinal cortex. J Comp Neurol, 391, 293–321.

Burwell, R. D., Bucci, D. J., Sanborn, M. R. & Jutras, M. J. 2004a. Perirhinal and postrhinal contributions to remote memory for context. J Neurosci, 24, 11023–8.

Burwell, R. D. & Hafeman, D. M. 2003. Positional firing properties of postrhinal cortex neurons. Neuroscience, 119, 577–88.

Burwell, R. D., Saddoris, M. P., Bucci, D. J. & Wiig, K. A. 2004b. Corticohippocampal contributions to spatial and contextual learning. J Neurosci, 24, 3826–36.

Dalton, M. A., Mccormick, C., De Luca, F., Clark, I. A. & Maguire, E. A. 2019. Functional connectivity along the anterior-posterior axis of hippocampal subfields in the ageing human brain. Hippocampus, 29, 1049–1062.

Diana, R. A. 2017. Parahippocampal Cortex Processes the Nonspatial Context of an Event. Cereb Cortex, 27, 1808–1816.

Doan, T. P., Lagartos-Donate, M. J., Nilssen, E. S., Ohara, S. & Witter, M. P. 2019. Convergent Projections from Perirhinal and Postrhinal Cortices Suggest a Multisensory Nature of Lateral, but Not Medial, Entorhinal Cortex. Cell Rep, 29, 617–627 e7.

Eichenbaum, H., Sauvage, M., Fortin, N., Komorowski, R. & Lipton, P. 2012. Towards a functional organization of episodic memory in the medial temporal lobe. Neurosci Biobehav Rev, 36, 1597–608.

Estela-Pro, V. J. & Burwell, R. D. 2022. The anatomy and function of the postrhinal cortex. Behav Neurosci, 136, 101–113.

Faivre, N., Dubois, J., Schwartz, N. & Mudrik, L. 2019. Imaging object-scene relations processing in visible and invisible natural scenes. Sci Rep, 9, 4567.

Fraser, K. M. & Holland, P. C. 2019. Occasion setting. Behav Neurosci, 133, 145–175.

Furtak, S. C., Ahmed, O. J. & Burwell, R. D. 2012. Single neuron activity and theta modulation in postrhinal cortex during visual object discrimination. Neuron, 76, 976–88.

Furtak, S. C., Cho, C. E., Kerr, K. M., Barredo, J. L., Alleyne, J. E., Patterson, Y. R. & Burwell, R. D. 2009. The Floor Projection Maze: A novel behavioral apparatus for presenting visual stimuli to rats. J Neurosci Methods, 181, 82–8.

Gaffan, E. A., Healey, A. N. & Eacott, M. J. 2004. Objects and positions in visual scenes: effects of perirhinal and postrhinal cortex lesions in the rat. Behav Neurosci, 118, 992–1010.

Gofman, X., Tocker, G., Weiss, S., Boccara, C. N., Lu, L., Moser, M. B., Moser, E. I., Morris, G. & Derdikman, D. 2019. Dissociation between Postrhinal Cortex and Downstream Parahippocampal Regions in the Representation of Egocentric Boundaries. Curr Biol, 29, 2751–2757.e4.

Heimer-Mcginn, V. R., Poeta, D. L., Aghi, K., Udawatta, M. & Burwell, R. D. 2017. Disconnection of the Perirhinal and Postrhinal Cortices Impairs Recognition of Objects in Context But Not Contextual Fear Conditioning. J Neurosci, 37, 4819–4829.

Ho, J. W. & Burwell, R. D. 2014. Perirhinal and Postrhinal Functional Inputs to the Hippocampus. In: Derdikman, D. & Knierim, J. J. (eds.) Space,Time and Memory in the Hippocampal Formation. Vienna: Springer Vienna.

Hwang, E., Willis, B. S. & Burwell, R. D. 2018. Prefrontal connections of the perirhinal and postrhinal cortices in the rat. Behav Brain Res, 354, 8–21.

Jacobson, T. K., Ho, J. W., Kent, B. W., Yang, F. C. & Burwell, R. D. 2014. Automated visual cognitive tasks for recording neural activity using a floor projection maze. J Vis Exp, e51316.

Karanian, J. M. & Slotnick, S. D. 2014. False memory for context activates the parahippocampal cortex. Cogn Neurosci, 5, 186–92.

Karanian, J. M. & Slotnick, S. D. 2017. False memory for context and true memory for context similarly activate the parahippocampal cortex. Cortex, 91, 79–88.

Komorowski, R. W., Manns, J. R. & Eichenbaum, H. 2009. Robust conjunctive item-place coding by hippocampal neurons parallels learning what happens where. J Neurosci, 29, 9918–29.

Kondo, H. & Witter, M. P. 2014. Topographic organization of orbitofrontal projections to the parahippocampal region in rats. J Comp Neurol, 522, 772–93.

Labar, K. S., Gitelman, D. R., Parrish, T. B., Kim, Y. H., Nobre, A. C. & Mesulam, M. M. 2001. Hunger selectively modulates corticolimbic activation to food stimuli in humans. Behav Neurosci, 115, 493–500.

Lachance, P. A., Graham, J., Shapiro, B. L., Morris, A. J. & Taube, J. S. 2022. Landmark-modulated directional coding in postrhinal cortex. Sci Adv, 8, eabg8404.

Lachance, P. A., Todd, T. P. & Taube, J. S. 2019. A sense of space in postrhinal cortex. Science, 365.

Lashley, K. S. 1930. The mechanism of vision: III. The comparative visual acuity of pigmented and albino rats. Journal of Genetic Psychology, 37, 481–484.

Lim, H. Y., Ahn, J. R. & Lee, I. 2022. The Interaction of Cue Type and Its Associated Behavioral Response Dissociates the Neural Activity between the Perirhinal and Postrhinal Cortices. eNeuro, 9.

Malkova, L. & Mishkin, M. 2003. One-trial memory for object-place associations after separate lesions of hippocampus and posterior parahippocampal region in the monkey. J Neurosci, 23, 1956–65.

Martin, C. B., Mclean, D. A., O’neil, E. B. & Kohler, S. 2013. Distinct familiarity-based response patterns for faces and buildings in perirhinal and parahippocampal cortex. J Neurosci, 33, 10915–23.

Mckenzie, S., Frank, A. J., Kinsky, N. R., Porter, B., Riviere, P. D. & Eichenbaum, H. 2014. Hippocampal representation of related and opposing memories develop within distinct, hierarchically organized neural schemas. Neuron, 83, 202–15.

Mullally, S. L. & Maguire, E. A. 2011. A new role for the parahippocampal cortex in representing space. J Neurosci, 31, 7441–9.

Nelson, M. J., Pouget, P., Nilsen, E. A., Patten, C. D. & Schall, J. D. 2008. Review of signal distortion through metal microelectrode recording circuits and filters. Journal of Neuroscience Methods, 169, 141–157.

Nishio, N., Tsukano, H., Hishida, R., Abe, M., Nakai, J., Kawamura, M., Aiba, A., Sakimura, K. & Shibuki, K. 2018. Higher visual responses in the temporal cortex of mice. Sci Rep, 8, 11136.

Norman, G. & Eacott, M. J. 2005. Dissociable effects of lesions to the perirhinal cortex and the postrhinal cortex on memory for context and objects in rats. Behav Neurosci, 119, 557–66.

Park, E. H., Ahn, J. R. & Lee, I. 2017. Interactions between stimulus and response types are more strongly represented in the entorhinal cortex than in its upstream regions in rats. Elife, 6.

Peng, X. & Burwell, R. D. 2021. Beyond the hippocampus: The role of parahippocampal-prefrontal communication in context-modulated behavior. Neurobiol Learn Mem, 185, 107520.

Ploner, C. J., Gaymard, B. M., Rivaud-Pechoux, S., Baulac, M., Clemenceau, S., Samson, S. & Pierrot-Deseilligny, C. 2000. Lesions affecting the parahippocampal cortex yield spatial memory deficits in humans. Cereb Cortex, 10, 1211–6.

Powers, M. K. & Green, D. G. 1978. Single retinal ganglion cell responses in the dar-reared rat: Grating acuity, contrast sensitivity, and defocusing. Vision Res, 18, 1533–1539.

Prusky, G. T., Harker, K. T., Douglas, R. M. & Whishaw, I. Q. 2002. Variation in visual acuity within pigmented, and between pigmented and albino rat strains. Behav Brain Res, 136, 339–48.

Qi, X., Du, Z. J., Zhu, L., Liu, X., Xu, H., Zhou, Z., Zhong, C., Li, S., Wang, L. & Zhang, Z. 2019. The Glutamatergic Postrhinal Cortex-Ventrolateral Orbitofrontal Cortex Pathway Regulates Spatial Memory Retrieval. Neurosci Bull, 35, 447–460.

Rémy, F., Vayssière, N., Saint-Aubert, L., Bacon-Macé, N., Pariente, J., Barbeau, E. & Fabre-Thorpe, M. 2020. Age effects on the neural processing of object-context associations in briefly flashed natural scenes. Neuropsychologia, 136, 107264.

Shields, G. S., Mccullough, A. M., Ritchey, M., Ranganath, C. & Yonelinas, A. P. 2019. Stress and the medial temporal lobe at rest: Functional connectivity is associated with both memory and cortisol. Psychoneuroendocrinology, 106, 138–146.

Stevenson, R. F., Reagh, Z. M., Chun, A. P., Murray, E. A. & Yassa, M. A. 2020. Pattern Separation and Source Memory Engage Distinct Hippocampal and Neocortical Regions during Retrieval. J Neurosci, 40, 843–851.

Suzuki, W. A. & Amaral, D. G. 1994. Topographic organization of the reciprocal connections between the monkey entorhinal cortex and the perirhinal and parahippocampal cortices. J Neurosci, 14, 1856–77.

Yeung, L. K., Olsen, R. K., Hong, B., Mihajlovic, V., D’angelo, M. C., Kacollja, A., Ryan, J. D. & Barense, M. D. 2019. Object-in-place Memory Predicted by Anterolateral Entorhinal Cortex and Parahippocampal Cortex Volume in Older Adults. J Cogn Neurosci, 31, 711–729.

Zhang, G. R., Zhao, H., Choi, E. M., Svestka, M., Wang, X., Nagayach, A., Singh, A., Cook, R. G. & Geller, A. I. 2019. An identified ensemble within a neocortical circuit encodes essential information for genetically-enhanced visual shape learning. Hippocampus, 29, 710–725.

Zhang, G. R., Zhao, H., Cook, N., Svestka, M., Choi, E. M., Jan, M., Cook, R. G. & Geller, A. I. 2017. Characteristic and intermingled neocortical circuits encode different visual object discriminations. Behav Brain Res, 331, 261–275.

Zhu, L., Wang, Z., Du, Z., Qi, X., Shu, H., Liu, D., Su, F., Ye, Q., Liu, X., Zhou, Z., Tang, Y., Song, R., Wang, X., Lin, L., Li, S., Han, Y., Wang, L. & Zhang, Z. 2020. Impaired Parahippocampal Gyrus-Orbitofrontal Cortex Circuit Associated with Visuospatial Memory Deficit as a Potential Biomarker and Interventional Approach for Alzheimer Disease. Neurosci Bull.

